# Phosphorylation and DNA Damage Resolution Coordinate SOX2-Mediated Reprogramming in vivo

**DOI:** 10.1101/2025.07.15.664998

**Authors:** Xiaoling Zhong, Yuhua Zou, Chun-Li Zhang

**Affiliations:** Department of Molecular Biology, University of Texas Southwestern Medical Center, Dallas, TX 75390, USA; Hamon Center for Regenerative Science and Medicine, University of Texas Southwestern Medical Center, Dallas, TX 75390, USA; O’Donnell Jr. Brain Institute, University of Texas Southwestern Medical Center, Dallas, TX 75390, USA

**Keywords:** SOX2, in vivo reprogramming, glia-to-neuron reprogramming, S251 phosphorylation, PRKDC, NHEJ, KU80, LIG4, p53

## Abstract

The stem cell factor SOX2 can reprogram resident glial cells into neurons in the adult mammalian central nervous system (CNS), but the molecular mechanisms underlying this process remain poorly understood. Here, we show that both SOX2 phosphorylation and the PRKDC-dependent non-homologous end joining (NHEJ) pathway are essential for SOX2-mediated in vivo glia-to-neuron reprogramming. A phospho-mimetic SOX2 mutant significantly enhances reprogramming efficiency without altering neuronal fate. Conversely, loss of PRKDC or knockdown of core NHEJ components KU80 and LIG4 abolishes reprogramming. Notably, p53 knockdown restores reprogramming in PRKDC-deficient mice. These findings demonstrate that SOX2-driven glial reprogramming requires both precise posttranslational regulation and effective DNA damage repair, and suggest that targeting these pathways could enhance regenerative strategies in the CNS.

## INTRODUCTION

With the exception of a few discrete neurogenic niches, the adult mammalian CNS has largely lost its capacity to generate new neurons for repair following injury or disease. One emerging strategy for overcoming such regeneration failure is through cell fate reprogramming in vivo ^1-4^. This strategy harnesses the abundance and intrinsic plasticity of endogenous glial cells and reprograms them into neurons by manipulating key fate-determining transcription factors and signaling pathways.

SOX2 is one of the most effective transcription factors for in vivo glial reprogramming. Its overexpression is sufficient to induce the generation of new neurons in the adult mouse brain and spinal cord ^5-10^. Rigorous genetic lineage tracing has demonstrated that these SOX2-induced neurons originate from resident glial cells including astrocytes and NG2 glia ^5,8^. Notably, SOX2-mediated glial reprogramming proceeds through a defined multi-step process, involving transient ASCL1^+^ progenitors and DCX^+^ neuroblasts, and culminating in the formation of NeuN^+^ mature neurons—particularly in the presence of neurotrophic factors such as BDNF and NOG ^5,6,10^.

The activity of transcription factors is often modulated by post-translational modifications such as phosphorylation. For example, the neurogenic potential of ASCL1 and NEUROG2 is negatively regulated by multi-site phosphorylation, and phospho-deficient mutants of these factors exhibit enhanced capacity to reprogram astroglia into neurons ^11-14^. Similarly, SOX2 undergoes phosphorylation by kinases such as PRKDC, the catalytic subunit of DNA-dependent protein kinase (DNA-PK) ^15^. However, the functional significance of SOX2 phosphorylation in the context of glia-to-neuron reprogramming remains unknown.

Cellular reprogramming is inherently stressful and is accompanied by extensive epigenetic remodeling and transcriptional changes, which can lead to the accumulation of DNA damage. Efficient DNA repair is therefore essential for maintaining genomic integrity and enabling successful reprogramming ^16-20^. While homologous recombination repair predominates in proliferative cells such as stem cells ^21,22^, the PRKDC-dependent nonhomologous end joining (NHEJ) pathway serves as the principal DNA repair mechanism in terminally differentiated cells like astrocytes ^23^. Despite this, the role of NHEJ in glia-to-neuron reprogramming in vivo has not been investigated.

In this study, we first assessed how phosphorylation affects SOX2-mediated reprogramming of resident glia in the adult mouse striatum and then examined the involvement of the PRKDC-dependent NHEJ pathway. Our findings uncover a previously unrecognized essential role for both SOX2 phosphorylation and the NHEJ repair mechanism in regulating glia-to-neuron reprogramming in vivo.

## RESULTS

### Phosphorylation of SOX2 at S251 is critical for its reprogramming activity in vivo

Given that phosphorylation at serine 251 (S251) enhances SOX2 protein stability and transcriptional activity ^15,24^, we investigated whether this post-translational modification influences SOX2’s ability to reprogram astrocytes in vivo. To mimic constitutive phosphorylation, we generated a phospho-mimetic SOX2 mutant (SOX2^S251E^) by substituting S251 with glutamic acid via site-directed mutagenesis. Conversely, a non-phosphorylatable mutant (SOX2^S251A^) was created by replacing S251 with alanine. As previously described ^5^, lentiviruses expressing wild-type SOX2, SOX2^S251E^, SOX2^S251A^, or GFP control under the human *GFAP* (*hGFAP*) promoter were injected into the striata of adult C57BL/6J mice (Figure 1A). Immunohistochemistry performed 4 weeks post virus-injection (wpv) confirmed robust expression of GFP and SOX2 variants in the targeted region (Figure 1B). Neurogenesis was then assessed by quantifying doublecortin (DCX)-positive cells in the injected striata. As expected ^5^, when compared to the GFP control, wild-type SOX2 induced a substantial number of DCX^+^ new neurons (Figure 1B-C; p<0.0001). Interestingly, the SOX2^S251E^ mutant triggered a striking 2.99-fold increase in DCX^+^ cells relative to wild-type SOX2 (Figure 1B-C; p=0.0106), suggesting enhanced reprogramming efficiency. In contrast, DCX^+^ cells were rarely observed in the striatum injected with virus expressing SOX2^S251A^ (Figure 1B-C). These findings indicate that phosphorylation at S251 is essential for SOX2-mediated glial reprogramming and highlight a critical post-translational mechanism regulating its reprogramming activity in vivo.

**Figure 1.**
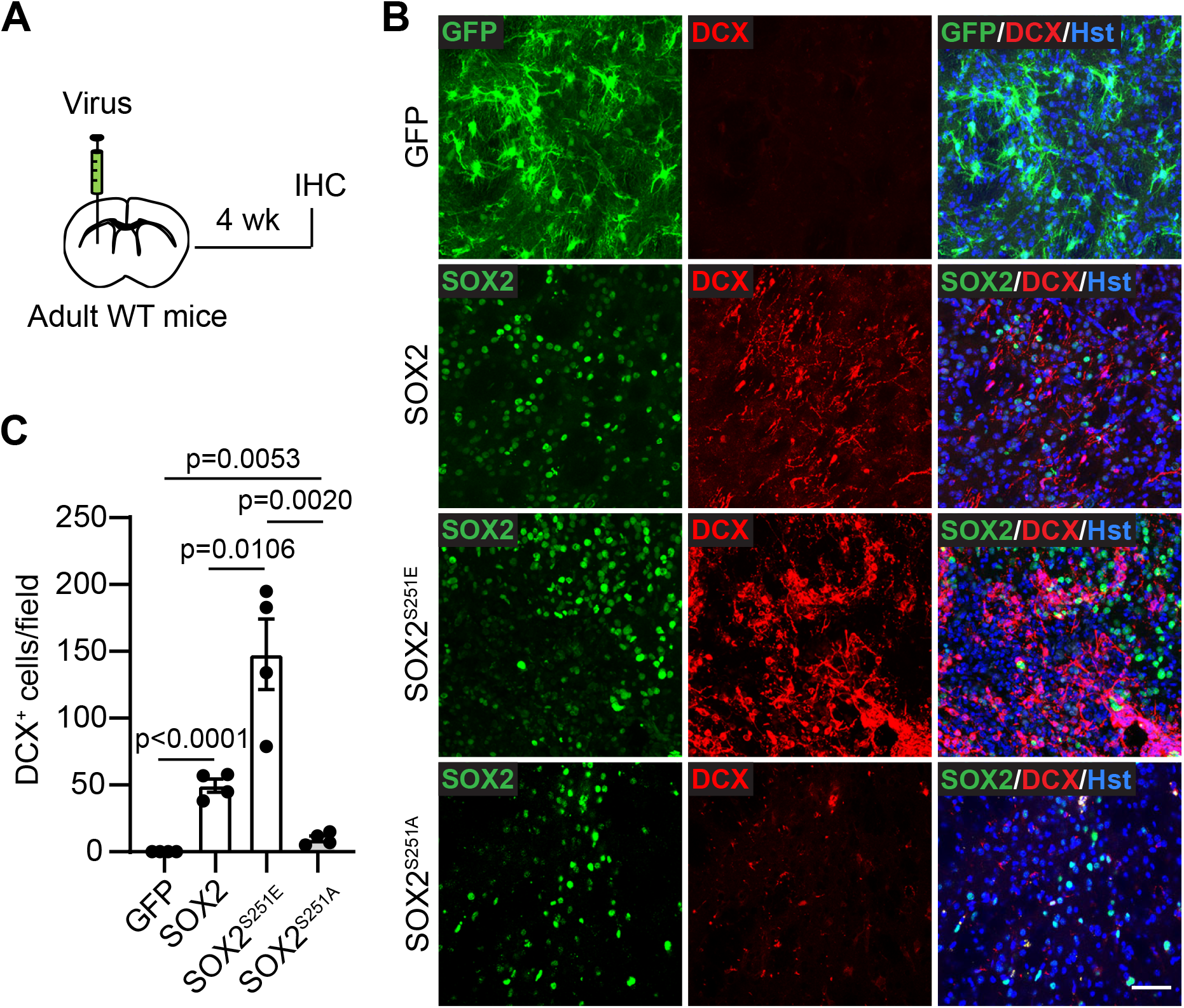
Phosphorylation of SOX2 at S251 is critical for its reprogramming activity in vivo. (A) Experimental design to determine the effect of SOX2 phosphorylation on reprogramming. Viruses were injected into the striatum of adult wild-type (WT) mice, and brains were analyzed by immunohistochemistry (IHC) 4 weeks (wk) later. (B) Representative confocal images showing expression of the indicated markers and DCX^+^ new neurons in the striatum surrounding the injection sites. Scale bar, 50 µm. (C)Quantification of DCX^+^ new neurons under the indicated conditions. n=4 mice for each group.

### SOX2 phosphorylation does not change the fate of reprogrammed cells

Since the phospho-mimetic mutant SOX2^S251E^ exhibited significantly enhanced reprogramming efficiency, we next asked whether it also altered the fate of the reprogrammed glia cells. To investigate this, we performed genetic lineage tracing in *Ascl1-CreER^T2^;R26R-tdTomato* (*tdT*) mice, as ASCL1^+^ neural progenitors are transiently induced during SOX2-mediated glial reprogramming in vivo ^6,7,9^. Adult mice received striatal injections of viruses expressing either wild-type SOX2 or the phospho-mimetic SOX2^S251E^, along with co-injection of BDNF and NOG to promote neuronal maturation ^5,6,9^. Tamoxifen was administered at 2 wpv, and brains were analyzed by immunohistochemistry at 10 wpv (Figure 2A).

**Figure 2.**
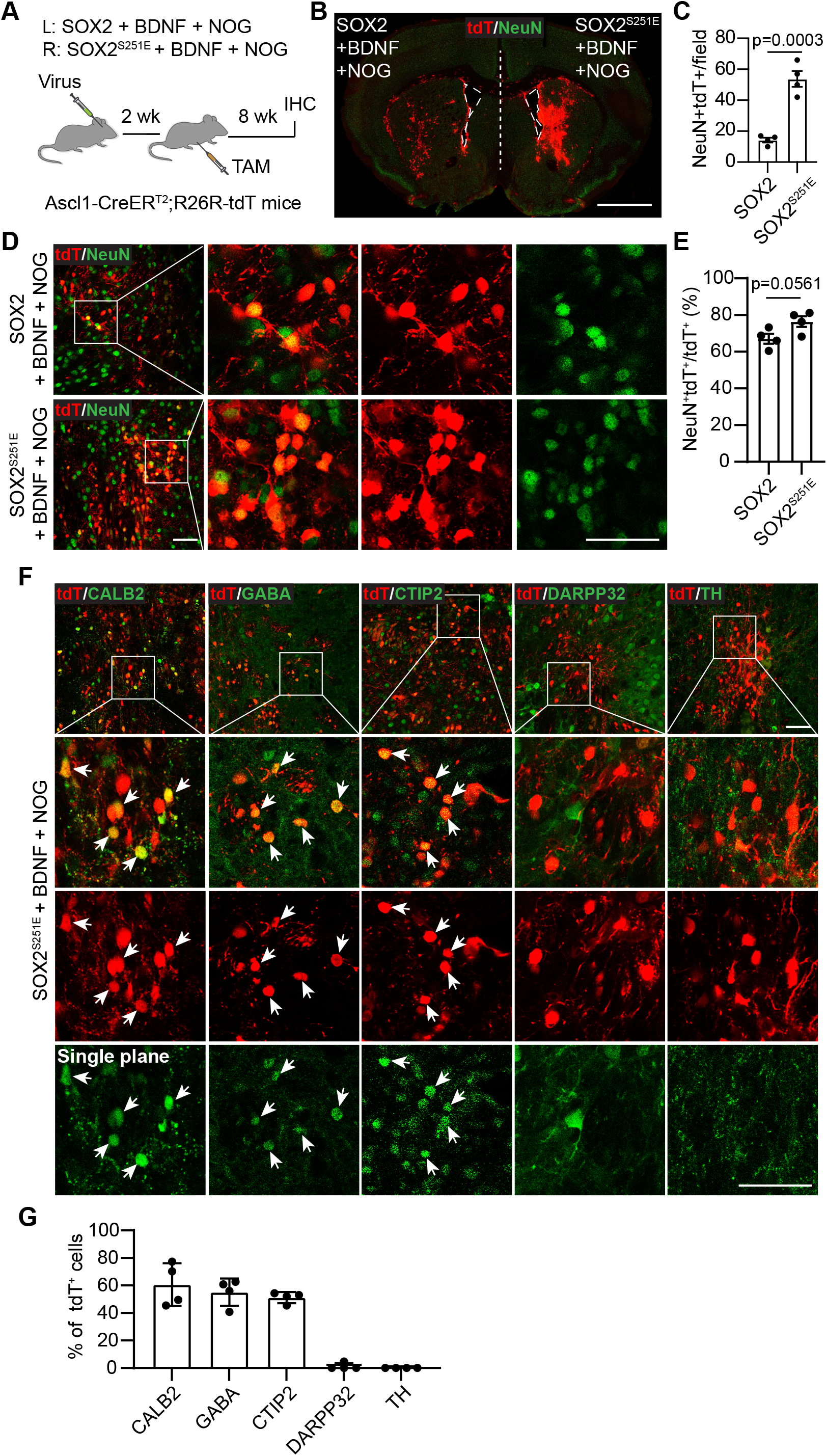
SOX2 phosphorylation does not change the fate of reprogrammed cells. (A) Experimental design to analyze the fate of reprogrammed cells. Mice were bilaterally injected with viruses into the striatum, treated with tamoxifen (TAM) 2 weeks (wk) later, and analyzed after another 8 weeks. (B) Low-magnification image of a brain section showing tdT-labeled reprogrammed cells. Scale bar, 1 mm. (C) Quantification of tdT-labeled neurons under the indicated conditions. n=4 mice for each group. (D) Representative confocal images showing tdT-labeled neurons surrounding the injection site. Scale bar, 50 µm. (E) Percentage of neurons among tdT-labeled cells. n=4 mice. (F) Representative confocal images showing neuronal subtypes of tdT-labeled cells. Scale bar, 50 µm. (G) Percentage distribution of neuronal subtypes among tdT-labeled cells. n=4 mice. See also Supplemental Figure S1.

Consistent with our earlier findings based on DCX^+^ cells, we observed a marked increase in tdT-labeled NeuN^+^ neurons in the striatum of mice injected with SOX2^S251E^ compared to those receiving wild-type SOX2 (Figure 2B-C; p = 0.0003). However, the proportion of tdT^+^ cells that became NeuN^+^ neurons remained similar between the two groups (67.11% for SOX2 and 76.43% for SOX2^S251E^; Figure 2D-E), suggesting that phosphorylation at S251 enhances overall reprogramming efficiency without altering the neuronal differentiation potential of reprogrammed astrocytes. To further assess neuronal subtype identity, we performed subtype marker analysis. Among tdT^+^NeuN^+^ cells in the SOX2^S251E^ group, approximately 60.66% expressed CALB2 (also known as calretinin), 55.12% were GABAergic, and 51.21% expressed CTIP2. In contrast, very few tdT^+^NeuN^+^ cells expressed DARPP32, and TH^+^ neurons were not detected (Figure 2F-G). These data indicate that while the majority of reprogrammed neurons adopt interneuron-like identities, few acquire projection neuron or dopaminergic fates. Interestingly, nearly 20% of tdT-labeled cells were NeuN^−^, indicating immaturation and/or alternative lineage differentiation. Further staining showed that the predominant NeuN^−^tdT^+^ population retained astrocytic identity, as marked by ALDOC and SOX9 (∼10%). The other glial lineage was OLIG2^+^ oligodendrocytes (Figure S1A-B). The distribution of these cell fates in the SOX2^S251E^ group was largely consistent with our previous observations using wild-type SOX2 ^6^.

Together, these results demonstrate that phosphorylation of SOX2 at S251 significantly enhances the in vivo reprogramming efficiency of astrocytes without altering their ultimate neuronal or glial fate choices.

### PRKDC is essential for SOX2-mediated in vivo reprogramming

Since PRKDC (also known as DNA-PKcs) is a major kinase responsible for phosphorylating SOX2 at S251 in glial cells ^15^, we investigated its role in SOX2-mediated glial reprogramming in vivo. Therefore, we injected SOX2-expressing viruses into the striata of adult wild-type or PRKDC-deficient (*Prkdc*^*scid*^) mice and performed immunohistochemical analysis at 4 wpv (Figure 3A). Robust SOX2 expression was observed in the striata of both wild-type and *Prkdc*^*scid*^ mice, indicating that PRKDC deficiency does not affect SOX2 expression per se (Figure 3B). As expected from our prior studies ^5,6^, SOX2 expression induced a substantial number of DCX^+^ cells in the striata of wild-type mice (Figure 3B-C). Strikingly, this reprogramming was completely abolished in *Prkdc*^*scid*^ mice (Figure 3B-C), indicating a critical requirement for PRKDC in SOX2-driven neurogenesis. To assess whether phosphorylation at S251 could bypass this requirement, we tested the phospho-mimetic SOX2^S251E^. However, SOX2^S251E^ also failed to induce DCX^+^ cells in the striata of *Prkdc*^*scid*^ mice (Figure 3B-C), suggesting that PRKDC may be required for SOX2-mediated reprogramming through additional mechanisms beyond S251 phosphorylation.

**Figure 3.**
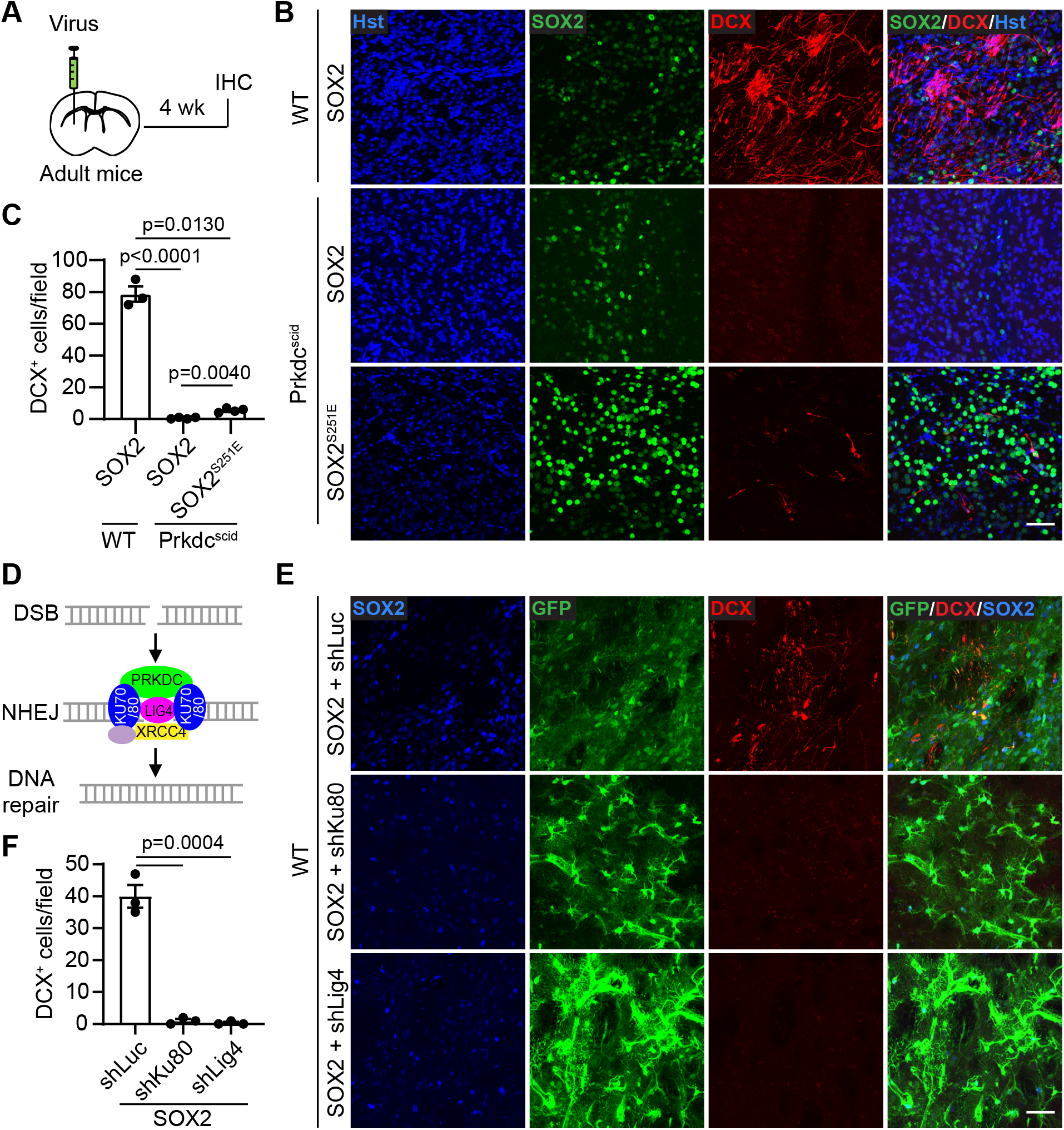
PRKDC-dependent NHEJ pathway is essential for SOX2-mediated in vivo reprogramming. (A) Experimental design to determine the effect of PRKDC deficiency on reprogramming. Adult wild-type (*WT*) or *Prkdc*^*scid*^ mice were injected with viruses and analyzed 4 weeks later. (B) Representative confocal images showing expression of the indicated markers surrounding the virus-injected striatal regions. Scale bar, 50 µm. (C) Quantification of DCX^+^ new neurons in the injected striatal regions. n=3 for *WT* mice and n=4 for *Prkdc*^*scid*^ mice per group. (D) Schematic diagram illustrating the NHEJ pathway for repairing DNA double-strand breaks. (E) Representative confocal images showing expression of the indicated markers surrounding the virus-injected striatal regions. Scale bar, 50 µm. (F) Quantification of DCX^+^ new neurons. n=3 mice for each group. See also Supplemental Figures S2 and S3.

To determine whether PRKDC deficiency broadly impairs adult neurogenesis, we examined endogenous DCX^+^ cells in two canonical neurogenic regions: the subventricular zone (SVZ) of the lateral ventricle and the subgranular zone (SGZ) of the dentate gyrus (DG). The distribution and abundance of DCX^+^ cells in these regions were comparable between wild-type and *Prkdc*^*scid*^ mice (Figure S2A-B), indicating that PRKDC is not essential for baseline adult neurogenesis. Together, these findings demonstrate that PRKDC is specifically required for SOX2-mediated glia-to-neuron reprogramming in vivo, but not for endogenous neurogenesis in the adult brain.

### The NHEJ pathway is critical for SOX2-mediated in vivo Reprogramming

PRKDC plays a central role in the NHEJ pathway for repairing DNA double-strand breaks, functioning both as the catalytic subunit and as a scaffold that recruits and connects core NHEJ components, including the KU70/80 heterodimer, LIG4, and XRCC4 ^25-27^ (Figure 3D). To determine whether other components of the NHEJ pathway are also required for SOX2-mediated in vivo reprogramming, we examined KU80 (also known as XRCC5) and LIG4 using an shRNA-mediated knockdown approach. Knockdown efficiency was validated in primary cultured astrocytes transduced with lentiviruses targeting *Ku80* or *Lig4*, with a luciferase-targeting shRNA (*shLuc*) used as a control (Figure S3). We then co-injected the most effective shRNAs together with SOX2-expressing virus into the striata of wild-type mice (Figure 3E-F). At 4 wpv, we analyzed SOX2 expression and the presence of DCX^+^ new neurons by immunostaining. Compared to the SOX2 + shLuc control group, which showed robust induction of DCX^+^ cells, knockdown of *Ku80* resulted in a near-complete loss of DCX^+^ cells (∼4–5 cells per field), indicating a severe disruption of reprogramming (Figure 3E-F). We also evaluated the impact of *Lig4* knockdown. Although the shRNA targeting *Lig4* produced only moderate transcript reduction (Figure S3B), it still remarkably reduced the number of DCX^+^ cells when co-expressed with SOX2 (Figure 3E-F), suggesting that even partial loss of LIG4 function impairs reprogramming efficiency. Together, these findings demonstrate that the NHEJ pathway is essential for SOX2-mediated neuronal reprogramming of astrocytes in vivo.

### SOX2-mediated reprogramming heightens DNA damage

The PRKDC-dependent NHEJ pathway is the major pathway for repairing DNA double-strand breaks in terminally differentiated astrocytes ^23^. Its requirement in SOX2-mediated reprogramming suggests that the reprogramming process may induce DNA damage, consistent with previous reports that cellular reprogramming is inherently stressful and leads to accumulated genomic instability ^20,28^. To test this hypothesis, we examined the expression of γH2AX, the phosphorylated form of histone H2AX, which serves as one of the earliest and most sensitive markers of DNA double-strand breaks ^29^. Because wild-type mice efficiently repair DNA damage, we analyzed γH2AX expression in *Prkdc*^*scid*^ mice, which are deficient in NHEJ-mediated repair. One week after striatal injection of lentiviruses expressing SOX2 or a GFP control, we observed a significant increase in the number of γH2AX foci in SOX2-expressing astrocytes compared to GFP controls (Figure 4A-C). These results indicate that SOX2-mediated reprogramming induces DNA double-strand breaks and highlight the critical role of PRKDC and the NHEJ pathway in resolving genomic stress during cellular reprogramming.

**Figure 4.**
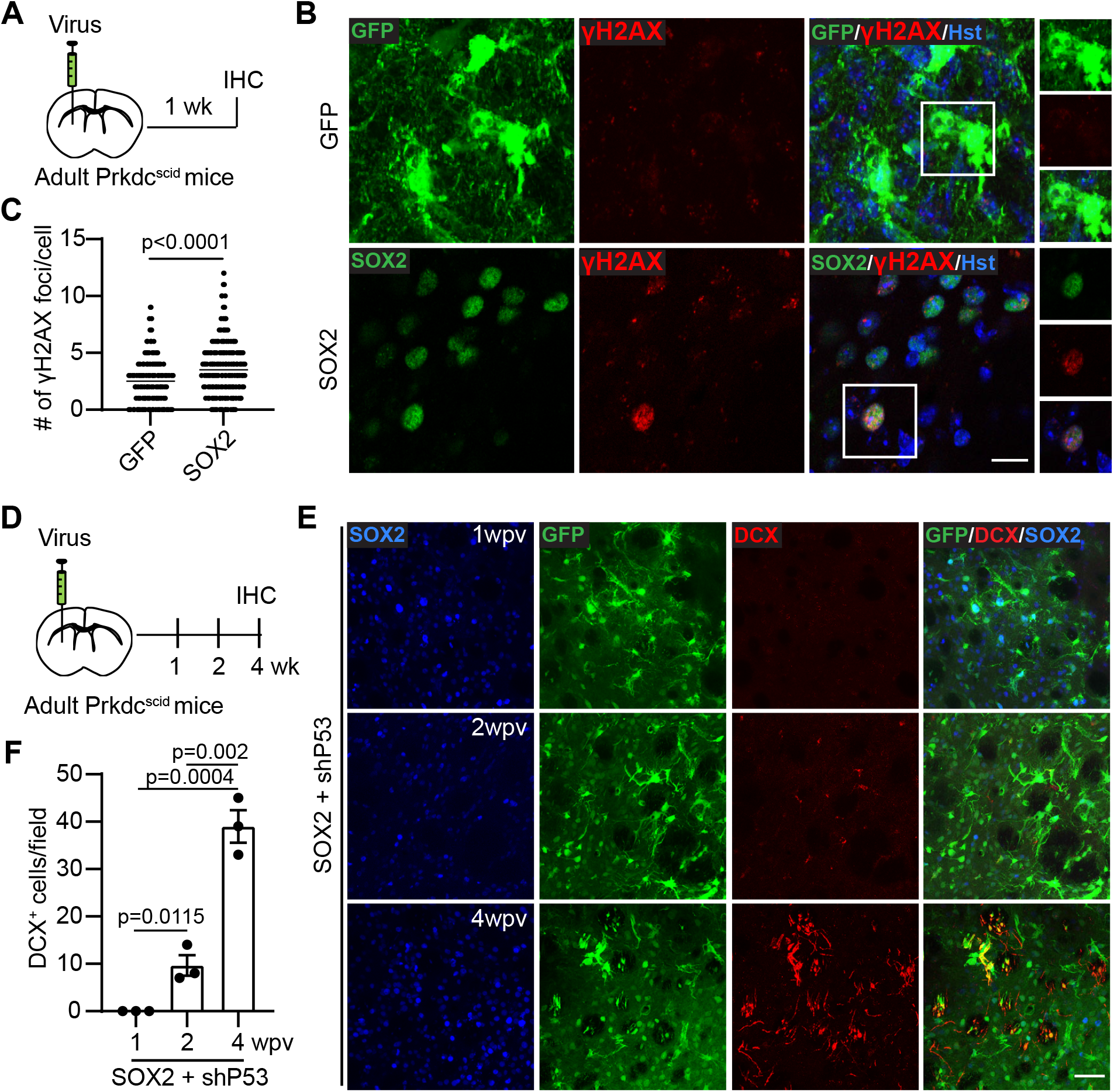
p53 knockdown rescues SOX2-mediated reprogramming in PRKDC-deficient mice. (A) Experimental design to determine the effect of SOX2-medaited reprogramming on DNA damage. (B) Representative confocal images showing expression of the indicated markers surrounding the virus-injected striatal regions. (C) Number of γH2AX foci in virus-transduced cells. n= 271 GFP^+^ cells from 3 mice and n=563 SOX2^+^ cells from 3 mice. (D) Experimental design to determine the effect of p53 knockdown on SOX2-mediated reprogramming in *Prkdc*^*scid*^ mice. (E) Representative confocal images showing expression of the indicated markers surrounding the virus-injected striatal regions. wpv, weeks post virus-injection. Scale bar, 50 µm. (F) Quantification of DCX^+^ new neurons. n=3 mice for each group. wpv, weeks post virus-injection.

### p53 knockdown rescues SOX2-mediated reprogramming in PRKDC-deficient mice

Given that disruption of the NHEJ pathway can lead to the accumulation of DNA double-strand breaks and trigger p53-dependent apoptotic responses ^30,31^, we hypothesized that excessive activation of p53 may serve as a key barrier to SOX2-mediated reprogramming in PRKDC-deficient astrocytes. To test this, we employed a previously validated *shRNA* construct known to efficiently knock down *p53* expression in vivo ^9^. Lentiviruses co-expressing *SOX2* and *shP53* were stereotactically injected into the striatum of *Prkdc*^*scid*^ mice, and the extent of reprogramming was assessed over a four-week time course (Figure 4D). Expression of both SOX2 and the shRNA (marked by co-expressed GFP) was confirmed in the targeted striatal regions (Figure 4E). DCX-positive new neurons were virtually undetectable at 1 wpv, with only sparse emergence at 2 wpv. However, by 4 wpv, a substantial number of DCX^+^ cells were observed, indicating robust reprogramming activity (Figure 4E-F). These findings suggest that suppressing *p53* alleviates a major apoptotic checkpoint that otherwise limits reprogramming efficiency in the context of impaired DNA repair. Together, this supports a model in which SOX2-driven reprogramming induces DNA damage that, in the absence of functional NHEJ, activates p53 to trigger cell death. Attenuation of p53 activity therefore allows reprogrammed cells to survive the initial stress and proceed through becoming a neuronal fate.

## DISCUSSION

Understanding the molecular mechanisms by which glial cells can be reprogrammed into neurons is essential for developing effective regeneration-based therapies. Our results demonstrate that both protein phosphorylation and the NHEJ DNA repair pathway are critical for SOX2-mediated reprogramming of resident glia into neurons in the adult mouse brain.

In contrast to ASCL1 and NEUROG2, whose neurogenic activity is inhibited by multi-site phosphorylation ^11-14^, phosphorylation of SOX2 at S251 enhances and is essential for its reprogramming capacity. This modification appears to protect SOX2 from ubiquitination and proteasomal degradation ^15,32^, while also promoting its nuclear localization and transcriptional activity ^24^. Consistent with this, we observed stronger SOX2 immunostaining in tissues expressing the phospho-mimetic SOX2^S251E^ mutant compared to wild-type SOX2 or the phospho-deficient SOX2^S251A^ mutant, despite equivalent viral titers being injected into the striatum. In addition to PRKDC ^15^, S251 can also be phosphorylated by other kinases, including MAPK p38 and GSK3β ^15,32^. The involvement of multiple kinases may account for the observation that SOX2-mediated reprogramming is rescued by p53 knockdown in PRKDC-deficient mice.

In addition to functioning as a SOX2 kinase, PRKDC is the catalytic subunit of the DNA-PK holoenzyme, which also includes the KU70/80 heterodimer and is essential for the NHEJ pathway that repairs DNA double-strand breaks ^33^. Using both the *Prkdc*^*scid*^ genetic mouse model and shRNA-mediated knockdown, we demonstrate that the PRKDC-dependent NHEJ pathway is critical for SOX2-mediated in vivo glial reprogramming. This requirement is further supported by knockdown of LIG4, the enzyme responsible for ligating DNA ends in the final step of the NHEJ pathway ^34^. The necessity of NHEJ in this context aligns with the idea that cellular reprogramming entails extensive epigenetic and transcriptional remodeling, processes often associated with DNA damage ^16-20^. Supporting this, we observed elevated staining of γH2AX, a well-established and sensitive marker of DNA double-strand breaks ^29^, in SOX2-expressing cells during the early stages of reprogramming.

Consistent with previous studies showing that *p53* deletion can rescue the embryonic lethality associated with NHEJ deficiency ^30,31^, our results demonstrate that *p53* knockdown restores SOX2-mediated glial reprogramming in *Prkdc*-deficient mice. While the precise mechanism remains unclear, one likely explanation is that *p53* suppression reduces apoptosis triggered by DNA double-strand breaks via the *p53*-dependent pathway ^35^. Furthermore, because *p53* negatively regulates homologous recombination repair ^36^, its loss may facilitate a compensatory upregulation of this high-fidelity, though less efficient, DNA repair pathway in reprogramming glial cells. Thus, by alleviating DNA damage-induced apoptosis and enabling alternative DNA repair pathways, *p53* attenuation likely facilitates the survival and neuronal reprogramming of glial cells under genotoxic stress.

Together, our findings uncover key molecular mechanisms underlying in vivo glia-to-neuron reprogramming and suggest that targeting posttranslational modifications and DNA damage response pathways could enhance the efficacy of reprogramming-based regenerative therapies.

## EXPERIMENTAL PROCEDURES

### Animals

Wild-type C57BL/6J mice and the following genetically modified strains were obtained from The Jackson Laboratory: *Prkdc*^*scid*^ (stock #001913), *Ascl1-CreER^T2^* (stock #012882), and *R26R-tdTomato* (stock #007914). Both male and female mice aged 8 weeks or older were used in all experiments. Animals were housed in a temperature-controlled facility under a standard 12-hour light/dark cycle, with ad libitum access to food and water. All experimental procedures were conducted in accordance with protocols approved by the Institutional Animal Care and Use Committee (IACUC) at UT Southwestern.

### Tamoxifen administration

Tamoxifen (T5648, Sigma) was dissolved at a concentration of 40 mg/mL in a 9:1 (v/v) mixture of sesame oil and ethanol. Mice received intraperitoneal injections at a dose of 1 mg per 10 g of body weight once daily for 3–5 consecutive days.

### Virus preparation and intracranial injections

Lentiviral vectors *LV-hGFAP-GFP* (Addgene #183906) and *LV-hGFAP-SOX2* (Addgene #183907) were previously described ^5^, in which GFP or SOX2 expression is driven by the synthetic 681-bp *gfaABC1D* promoter ^37^. SOX2 mutant constructs were generated by Q5 site-directed mutagenesis kit (NEB, Cat No: E0554S). Lentiviral vectors *LV-hGFAP-GFP-shP53* and *LV-hGFAP-GFP-shLuc* were described previously ^9^; both GFP and miR30-based shRNAs are expressed under the control of the *hGFAP* promoter. Additional candidate shRNA sequences were subcloned into the *LV-hGFAP-GFP-shLuc* vector by replacing the *shLuc* sequence. The following shRNA sequences were used in this study: *shKu80 #1*: 5′-TTTCTTGTAATATATGTCCTTA-3′; *shKu80 #2*: 5′-TTATAAATTTGAATAGGGTCTA-3′; *shKu80 #3*: 5′-TTTATATTTCATTTGTTCCTCA-3′; *shLig4 #1*: 5′-TTTCAATAAAATTAATATCTTA-3′; *shLig4 #2*: 5′-TTTGATATCAAACTTGACCCCC-3′; *shLig4 #3*: 5′-TTTTCAATAAAATTAATATCTT-3′. All plasmids were verified by restriction enzyme digestion and Sanger sequencing. VSV-G pseudotyped lentiviruses were produced as previously described ^5,7,9^. For in vivo delivery, 3.0 µL of lentivirus (0.5-1 × 10^9^ pfu/mL) was injected stereotaxically into the striatum using a Hamilton syringe with a 34-gauge needle. The injection coordinates were: anterior/posterior, +1.0 mm; medial/lateral, ±2.0 mm; and dorsal/ventral from the skull surface, ™3.2 mm. For multifactor experiments, equal ratios of lentiviruses were mixed prior to injection. To minimize animal usage, bilateral injections with different vectors were performed when appropriate.

### Primary astrocyte culture and viral transduction

Primary astrocytes from postnatal mouse cerebral cortex were prepared following previously published protocols ^38,39^. Briefly, C57BL/6J mice at postnatal day 2–4 (P2–P4) were euthanized by decapitation, and brains were rapidly collected in ice-cold HBSS (GIBCO). After careful removal of the meninges, the gray matter of the cerebral cortex was dissected and mechanically dissociated. The resulting cell suspension was centrifuged at 400 × g for 5 minutes, resuspended, and plated in high-glucose DMEM supplemented with 10% fetal bovine serum (FBS). After 7–10 days in vitro, cells were detached using trypsin/EDTA (GIBCO) and replated onto plastic dishes in the same medium. To enrich for astrocytes, loosely attached microglia, macrophages, and oligodendrocyte precursor cells were removed by vigorous shaking or by blasting the culture surface with medium using a pipette. The majority of cells in the resulting cultures were positive for glial fibrillary acidic protein (GFAP), confirming astrocytic identity. Cultures were maintained for up to 3 weeks, with medium changes every 3 days. For all experiments, astrocyte cultures were randomly assigned to experimental groups and transduced with the indicated lentiviral vectors.

### RNA extraction and qRT-PCR

Total RNA was extracted using TRIzol reagent (Thermo Fisher Scientific) according to the manufacturer’s instructions, followed by treatment with DNase I (Thermo Fisher Scientific) to remove genomic DNA contamination. Reverse transcription was performed using the High-Capacity cDNA Reverse Transcription Kit (Thermo Fisher Scientific). Quantitative PCR (qPCR) was conducted at 60 °C using Power SYBR Green PCR Master Mix (Thermo Fisher Scientific) on a QuantStudio 5 Real-Time PCR System (Applied Biosystems).

### Tissue preparation and immunofluorescence staining

Mice were euthanized and perfused with ice-cold PBS and 4% paraformaldehyde (PFA) in PBS. Brains were isolated and post-fixed with 4% PFA overnight at 4 °C. They were then cryoprotected with 30% sucrose solution in PBS for 24 hours and cut into 40-μm thick coronal sections. Sections were serially collected and stored in an anti-freezing solution at -20°C. Immunostaining procedure was conducted as previously described ^5,40^. The following primary antibodies were used: rabbit anti-DCX (1:500; CST,4604), goat anti-DCX (1:500; Santa Cruz, 8066), chicken anti-GFP (1:1,000; Aves Labs), rabbit anti-NeuN (1:1000; abcam,177487), goat anti-SOX2 (1:200; Sant Cruz,17320), rabbit anti-SOX2 (1:400; Millipore, AB5603), mouse anti-γH2AX (1:200; CST, 80312), rabbit anti-Calretinin (1:100; Abcam, Ab702), rabbit anti-GABA (1:500; Sigma, A2052), rat anti-CTIP2 (1:400; Millipore, MABE1045), rabbit anti-DARPP32 (1:500; Abcam, 40801), chicken anti-TH (1:400; Aves), goat anti-ALDOC (1:50; Sant cruz,12065), goat anti-SOX9 (1:200; R&D, AF3075), rabbit anti-OLIG2 (1:400; Millipore, Ab9610), goat anti-OLIG2 (1:400; R&D, AF2418). Appropriate Alexa Fluor-conjugated secondary antibodies were purchased from Invitrogen. Cell nuclei were counterstained with Hoechst 33342 (Hst) when appropriate. Images were taken using a Zeiss LSM700 confocal microscope for analysis.

## Statistical Analysis

The quantification data were expressed as mean ± SEM from three to four mice. Statistical analysis was performed by the unpaired homoscedastic Student’s t tests where appropriate. Differences were considered statistically significant at p < 0.05.

## Materials availability

All materials generated in this study are available from the lead contact.

## ACKNOWLEDGMENTS

We thank members of the C.-L.Z. laboratory for helpful discussions, reagents and technical assistance; C.-L.Z. is a W.W. Caruth, Jr. Scholar in Biomedical Research and supported by the TARCC, the Decherd Foundation, and NIH grants NS127375, NS117065, NS131489, NS111776.

## AUTHOR CONTRIBUTIONS

X.Z. and C.-L.Z. conceived and designed the experiments; X.Z. performed research, created figures, and drafted the original manuscript; Y.Z. provided essential technical support. C.-L.Z revised the manuscript. All authors reviewed and approved the final manuscript.

