## Supplemental Figures for "Phosphorylation and DNA Damage Resolution Coordinate SOX2-Mediated Reprogramming in vivo"

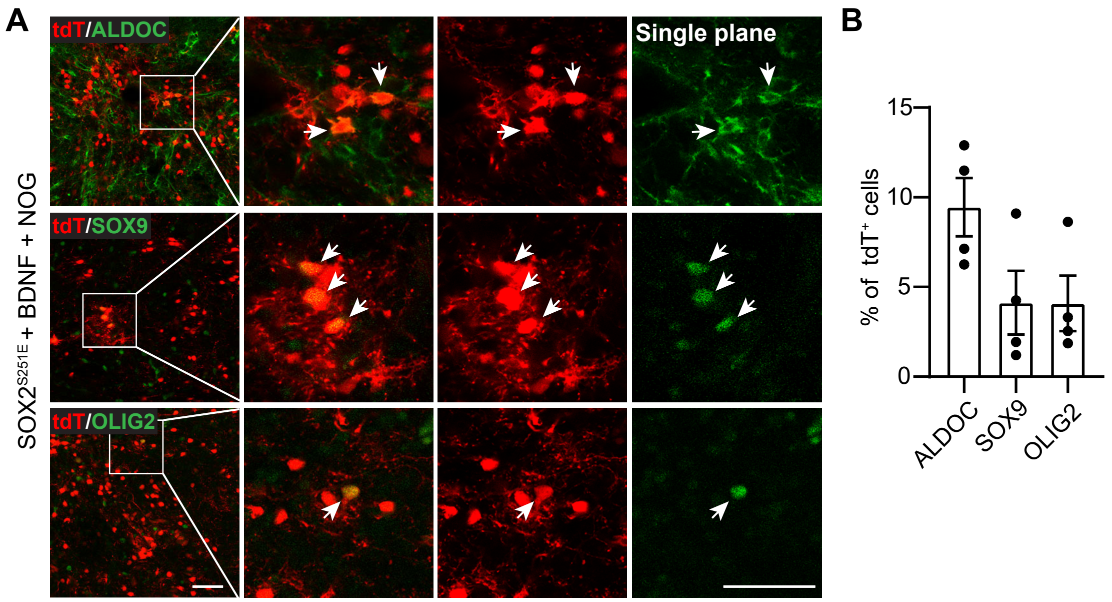


**Figure S1. Glial cell fate of SOX2$^{S251E}$-reprogrammed cells. Related to Figure 2.**
(A) Representative confocal images showing expression of the indicated markers surrounding the virus-injected striatal regions. Scale bar, 50 μm.
(B) Percentage distribution of glial cell types among tdT-labeled cells. n=4 mice.

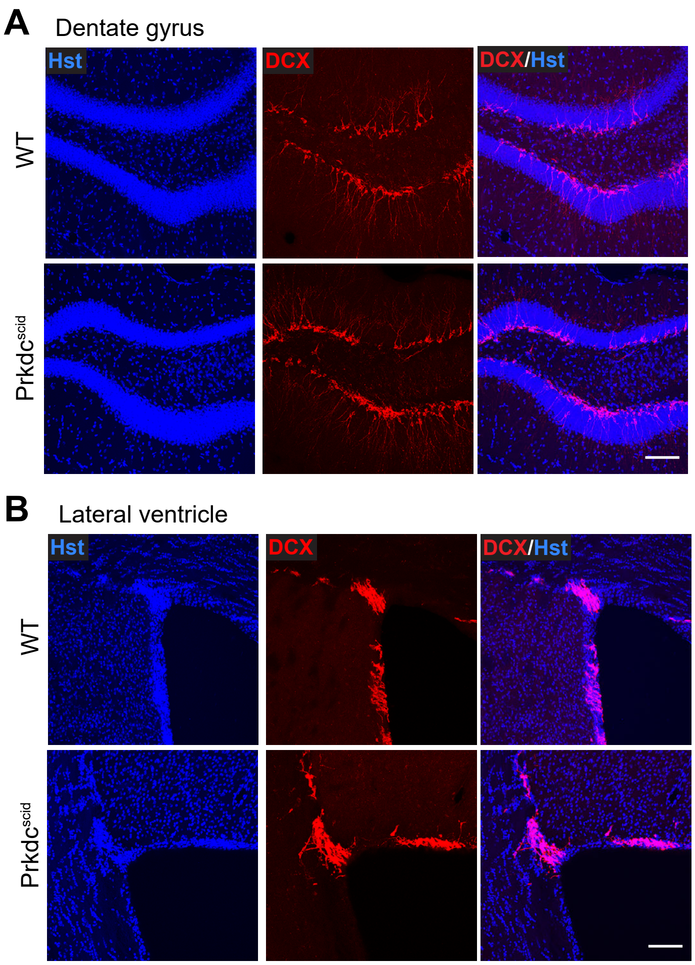


**Figure S2. PRKDC deficiency does not noticeably affect endogenous adult neurogenesis. Related to Figure 3.**

(A) Representative confocal images showing DCX$^{+}$ cells in the dentate gyrus. Scale bar, 50 µm.

(B) Representative confocal images showing DCX$^{+}$ cells in the lateral ventricle. Scale bar, 50 µm.

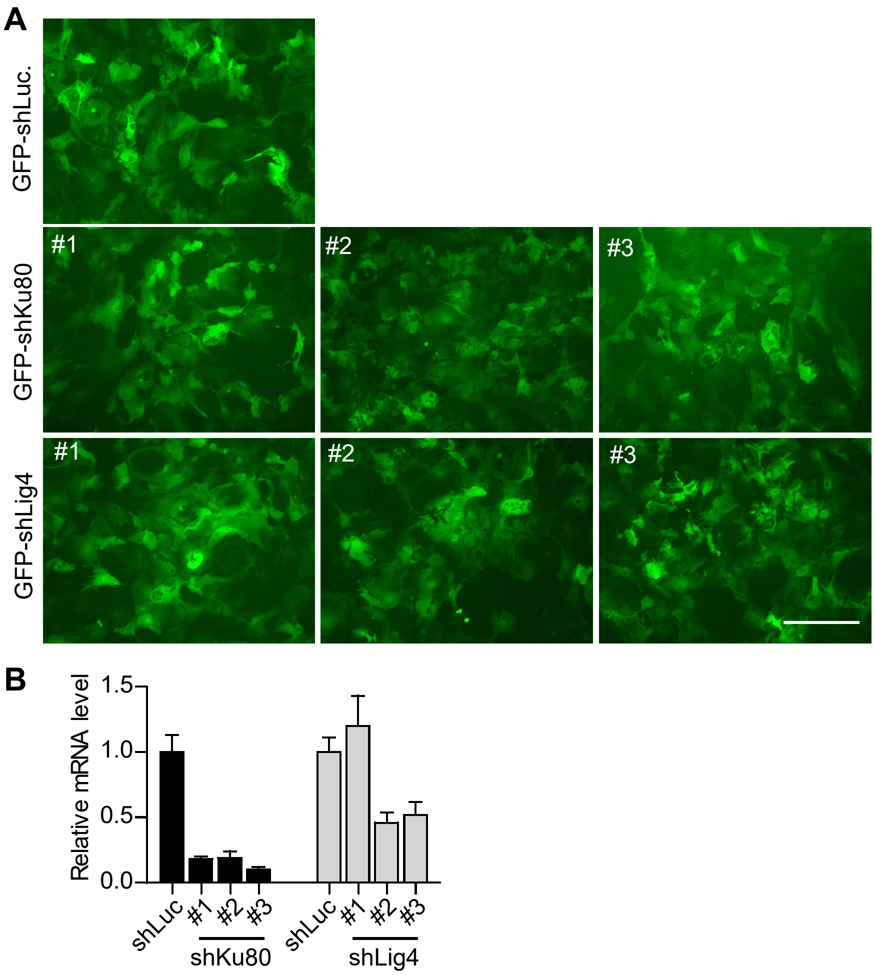


**Figure S3. Knockdown efficiency of shRNAs in primary astrocytes. Related to Figure 3.**
(A) Representative images of primary astrocytes transduced with shRNA-expressing viruses, indicated by co-expressed GFP. Scale bar, 400 μm.
(B) Knockdown efficiency of shRNAs was assessed by qRT-PCR. *shKu80-#3* and *shLig4-#2* were selected for in vivo experiments.
